# Human Milk Oligosaccharide 2’-fucosyllactose inhibits ligand binding to C-type lectin DC-SIGN but not to Langerin

**DOI:** 10.1101/2022.07.27.501236

**Authors:** Reshmi Mukherjee, Victor J. Somovilla, Fabrizio Chiodo, Sven Bruijns, Roland J Pieters, Johan Garssen, Yvette van Kooyk, Aletta D Kraneveld, Jeroen van Bergenhenegouwen

**Author notes:** shared last author (Aletta D Kraneveld, Jeroen van Bergenhenegouwen).

## Abstract

Human milk oligosaccharides (HMOs) and its most abundant component, 2’-Fucosyllactose (2’-FL), are known to be immunomodulatory. Previously, it was shown that HMOs and 2’-FL bind to the C-type lectin receptor DC-SIGN. Here we show, using a ligand-receptor competition assay, that a whole mixture of HMOs from pooled human milk (HMOS) and 2’-FL inhibit the binding of the carbohydrate-binding receptor DC-SIGN to its prototypical ligands, fucose and the oligosaccharide Lewis-B, (Le^b^) in a dose-dependent way. Interestingly, such inhibition by HMOS and 2’-FL was not detected for another C-type lectin, Langerin, evolutionary similar to DC-SIGN. The cell-ligand competition assay using DC-SIGN expressing cells confirmed that 2’-FL inhibits the binding of DC-SIGN to Le^b^. Molecular dynamics (MD) simulations show that 2’-FL exists in a preorganized bioactive conformation before binding to DC-SIGN and this conformation is retained after binding to DC-SIGN. Le^b^ has more flexible conformations and utilizes two binding modes, which operate one at a time via its two fucoses to bind to DC-SIGN. 2’-FL may have a reduced entropic penalty due to its preorganized state compared to Le^b^, and it has lower binding enthalpy, suggesting better binding to DC-SIGN. Thus, due to the better binding to DC-SIGN, 2’-FL may replace Le^b^ from its binding pocket in DC-SIGN. MD simulations also showed that 2’-FL does not bind to Langerin. Our studies confirm 2’-FL as a specific ligand for DC-SIGN and suggest that 2’-FL can replace other DC-SIGN ligands from its binding pocket during ligand-receptor interactions in possible immunomodulatory processes.

## Introduction

The C-type lectin receptors (CLR) family are pattern recognition receptors that play an important role in maintaining host tissue homeostasis but are also critically important in host-microbe interactions (Chiodo et al. 2021). The family of CLRs consists of many different subgroups harbouring several different receptors that all share a calcium-dependent carbohydrate-recognition domain (CRD) (Weis et al. 1998). CLRs respond to a wide variety of ligands both from endogenous and exogenous origins being (glyco)lipids, (glyco)proteins, or glycans (Foster et al. 2016).

Two of the best-studied CLRs are Dendritic Cell-Specific Intercellular Adhesion Molecule-3-Grabbing Non-integrin (DC-SIGN) and Langerin which recognize various glycan motifs (Valverde et al. 2020). DC-SIGN is expressed by immature dendritic cells from the dermis, mucosal tissues, and lymph nodes (van Vliet et al. 2008). Langerin is expressed by Langerhans cells, a subset of immature dendritic cells found in the epidermis of the human skin, lung, kidney, and liver (Merad et al. 2008). DC-SIGN and Langerin both have a glutamic acid-proline-asparagine (EPN) amino acid binding motif in their CRDs and use that motif to interact with fucose monosaccharide as well as fucose-containing motifs like Lewis-type antigens Lewis - X (Le^x^), Lewis-Y (Le^y^), Lewis-A (Le^a^), and Lewis-B (Le^b^). In most cases, DC-SIGN recognizes all four Lewis antigens (Valverde et al 2019; Pederson et al. 2014); however, Langerin recognizes mostly Le^b^ and Le^Y^ (Valverde et al. 2020; Lee et al. 2011).

Human milk oligosaccharides (HMOs) are abundant in human breast milk. HMOs contain mostly lactose (∼70g/L) and other various complex glycan structures (5-20 g/L) that are derived from lactose (Thurl et al. 2017; Urashima et al. 2012; Newburg DS. 2013). In the last decade interest in HMOs has surged and several health benefits have been attributed to them. HMOs are primarily seen as prebiotic substrates for beneficial bacteria, but accumulating research suggests that HMOs also have anti-pathogenic properties (Asadpoor et al. 2020). Moreover, HMOs have been found to directly interact with host cells promoting gut health by improving epithelial barrier function (Holscher et al. 2017; Holscher et al. 2014) but also by direct immunomodulating effects such as the activation or inhibition of Toll-like receptor (TLR) signalling (Cheng et al. 2019; Sodhi et al. 2021). However, the exact mechanism of action of direct immunomodulation by HMOs remains to be determined.

2’-fucosyllactose (2’-FL) is one of the most abundant HMOs and has also been shown to exert a direct effect on host immune cells or tissues as well as indirect effects by acting as a decoy receptor to inhibit the adhesion of pathogens (He et al. 2016; Goehring et al. 2016; Ruiz-Palacios et al. 2003; Weichert et al. 2013; Laucirica et al. 2017; Koromyslova et al. 2017). Currently, 2’-FL can be produced via different methods. In the past, 2’-FL was chemically synthesized from lactose, while recently, 2’-FL is biotechnologically produced on a large scale (Bych et al. 2019).

To explain the immunomodulatory functions of HMOs, researchers have investigated carbohydrate-binding receptors for HMOs binding. Using glycan-microarrays, it was shown that HMOs specifically interact with different lectins based on their structure and affinity (Noll et al. 2016). 2’-FL was shown to specifically interact with DC-SIGN, but not with other CLRs (Noll et al. 2016).

Here, we aim to study the effect of 2′-FL on the binding of DC-SIGN and Langerin to their prototypical ligands fucose monosaccharide and Le^b^ whose structure is depicted below (Fig 1). In addition, we investigated the effect of three different preparations of 2′-FL, biotechnologically produced commercially available 2′-FL (c-2′-FL), purified c-2′-FL (p2′-FL) and chemically synthesized 2′-FL (s2′-FL) on the binding of those prototypical ligands to DC-SIGN. Molecular dynamics (MD) simulations were run to understand the molecular details and dynamics of the interaction of 2′-FL with DC-SIGN and Langerin. In addition, MD simulation was performed for the interaction of Le^b^ with DC-SIGN to understand whether 2′-FL might be able to displace Le^b^ from the CRD of DC-SIGN.

**Figure 1.**
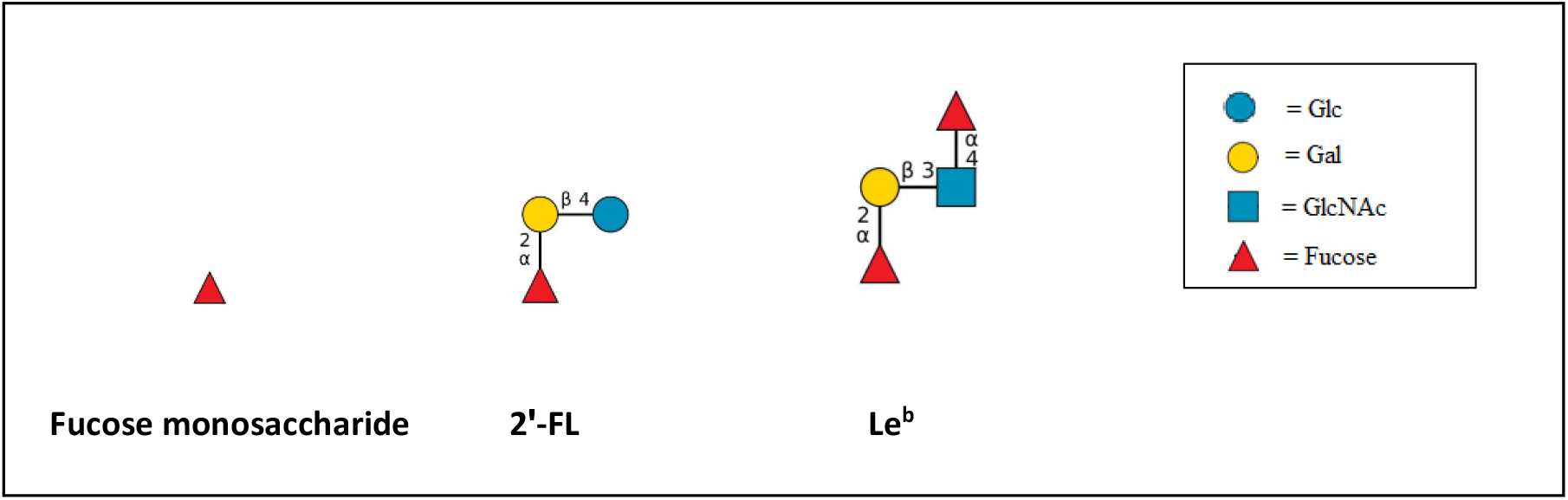
The schematic diagram of fucose monosaccharide, 2′-FL and the terminal tetrasaccharide of Le^b^ antigen are shown. Glc = glucose, Gal = galactose, GlcNAc = N-acetylglucosamine. The structures are drawn using DrawGlycan (Cheng et al. 2017).

## Results

### c2′-FL, p2′-FL, s2′-FL and HMOs inhibit the ligand binding to DC-SIGN, but not to Langerin

A ligand-receptor competition assay (solid-phase assay ELISA) was performed using commercially available multivalent polyacrylamide (PAA)-linked fucose monosaccharide (PAA-fucose) or PAA-linked-Le^b^ (PAA-Le^b^) as ligands. These were used as coating agents for these experiments. CLR-Fc constructs of DC-SIGN and Langerin were used as receptors. Maltotriose (MAL) was used as a negative control. EGTA binds to Ca^+2^ ion and thus it was used for inhibition of CLR-carbohydrate binding. To investigate whether the side-products in biotechnologically produced 2′-FL preparations (c2′-FL) play a role in affecting ligand-CLRs binding, we compared c2′-FL with purified c2′-FL (p2′-FL) and chemically synthesized 2′-FL (s2′-FL). As a binding control, a whole mixture of HMOs isolated from pooled human milk (HMOS) was used. c2′-FL, p2′-FL, s2′-FL and HMOS were used at a concentration of 1 µg/mL, 10 µg/mL, 100 µg/mL and 1000 µg/mL. c2′-FL, p2′-FL, s2′-FL and HMOS significantly inhibited the binding of DC-SIGN-Fc (used at 5 µg/mL) to the ligand, PAA-Fucose and PAA-Le^b^ in a dose-dependent way (Fig.2A and 2B). The inhibition occurred for all the 2′-FLs and HMOS only at 1000 µg/mL (1 mg/mL, 2 mM for 2′-FLs) concentration. On the contrary, c2′-FL, p2′-FL, s2′-FL and HMOS did not inhibit the binding of Langerin-Fc to the ligands PAA-fucose or PAA-Le^b^ (Fig 2C and 2D). When the assay was performed with a lower concentration of DC-SIGN-Fc (0.5 µg/mL and 0. 1 µg/mL), a similar effect on the binding of DC-SIGN by all the 2′-FLs and HMOS was observed (Fig. S1A-D).

**Figure 2.**
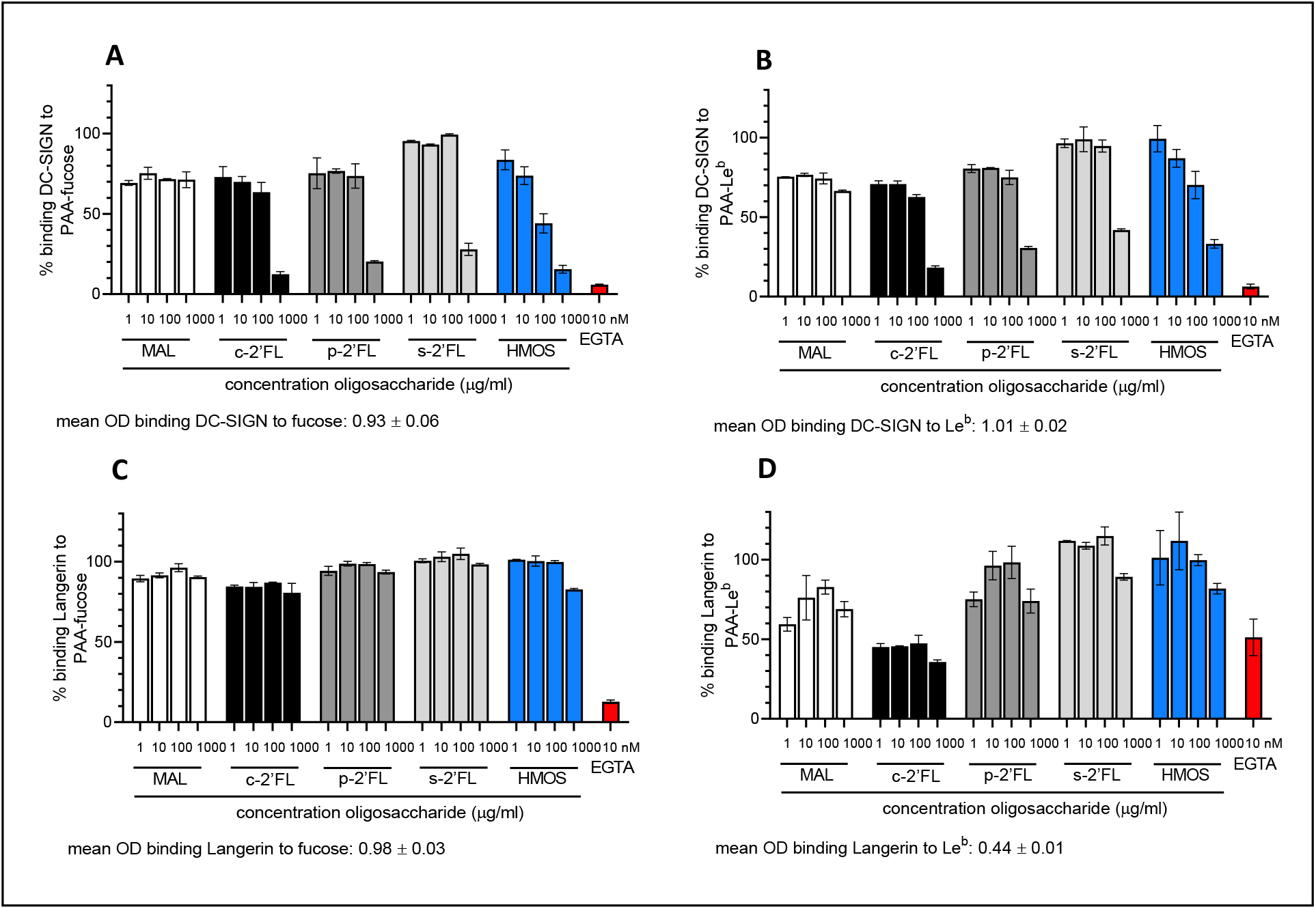
Ligand-receptor competition assay for DC-SIGN-Fc and Langerin-Fc (5 µg/mL) binding to PAA-fucose or PAA-Le^b^ with maltotriose (MAL, negative control), c2′-FL, p2′-FL, s2′-FL and HMOS [in various concentration of 1 µg/mL, 10 µg/mL, 100 µg/mL and 1000 µg/mL]. EGTA [10 nM] was used to demonstrate Ca^+2^-and Mg^+2^ dependent binding of DC-SIGN and Langerin to the PAA-fucose and PAA-Le^b^ ligands. A) inhibition by 2’-FLs and HMOS of the binding of DC-SIGN-Fc (5 µg/mL) to the ligand PAA-fucose; B) inhibition by 2’-FLs and HMOS of the binding of DC-SIGN-Fc (5 µg/mL) to the ligand PAA-Le^b^; C) No effect of 2’-FLs and HMOS on the binding of Langerin-Fc (5 µg/mL) to the ligand PAA-fucose; D) No effect of 2’-FLs and HMOS on the binding of Langerin-Fc (5 µg/mL) to the ligand PAA-Le^b^. Results are presented as mean % binding of ligand to DC-SIGN-Fc and Langerin-Fc ± SD, n=2 separate experiments. No block condition is regarded as 100% ligand-DC-SIGN-Fc/Langerin-Fc binding. c-2′-FL = biotechnologically produced commercially available 2′-FL, p2′-FL = purified c-2′-FL and s2′-FL = chemically synthesized 2′-FL.

### c2′-FL, p2′-FL, and s2′-FL inhibit the binding of Le^b^ to DC-SIGN expressing cells

A cell-ligand competition assay was performed by using DC-SIGN expressing OUW cells using Le^b^-coated fluorescent beads as a multivalent bona fide ligand. Anti-DC-SIGN mAbs AZN-D1 was used as a positive control to block the cellular interactions between DC-SIGN and Le^b^-coated fluorescent beads. Isotype antibody was used as a negative control to differentiate non-specific background signals. The DC-SIGN specific blocking antibody, AZN-D1, as well as EGTA inhibited the binding of Le^b^-coated fluorescent beads to DC-SIGN expressing cells. These results indicate specific binding of Le^b^ to DC-SIGN on the DC-SIGN-expressing cells. The DC-SIGN expressing cells binding to Le^b^ were inhibited by c2′-FL, p2′-FL, s2′-FL, and HMOS at a concentration of 5 mg/mL (Fig 3). Here, we observe again that 2′-FL can mimic the inhibitory effect of the HMOs which is a mixture of various complex oligosaccharides. But this effect was not observed at 1 mg/mL concentration of all 2′-FLs (Fig. 3 insert). All 2′-FLs lost the efficiency of inhibition of binding of Le^b^ to DC-SIGN expressing cells, but the HMOS (1 mg/mL) retained some of the inhibition at this concentration.

**Figure 3.**
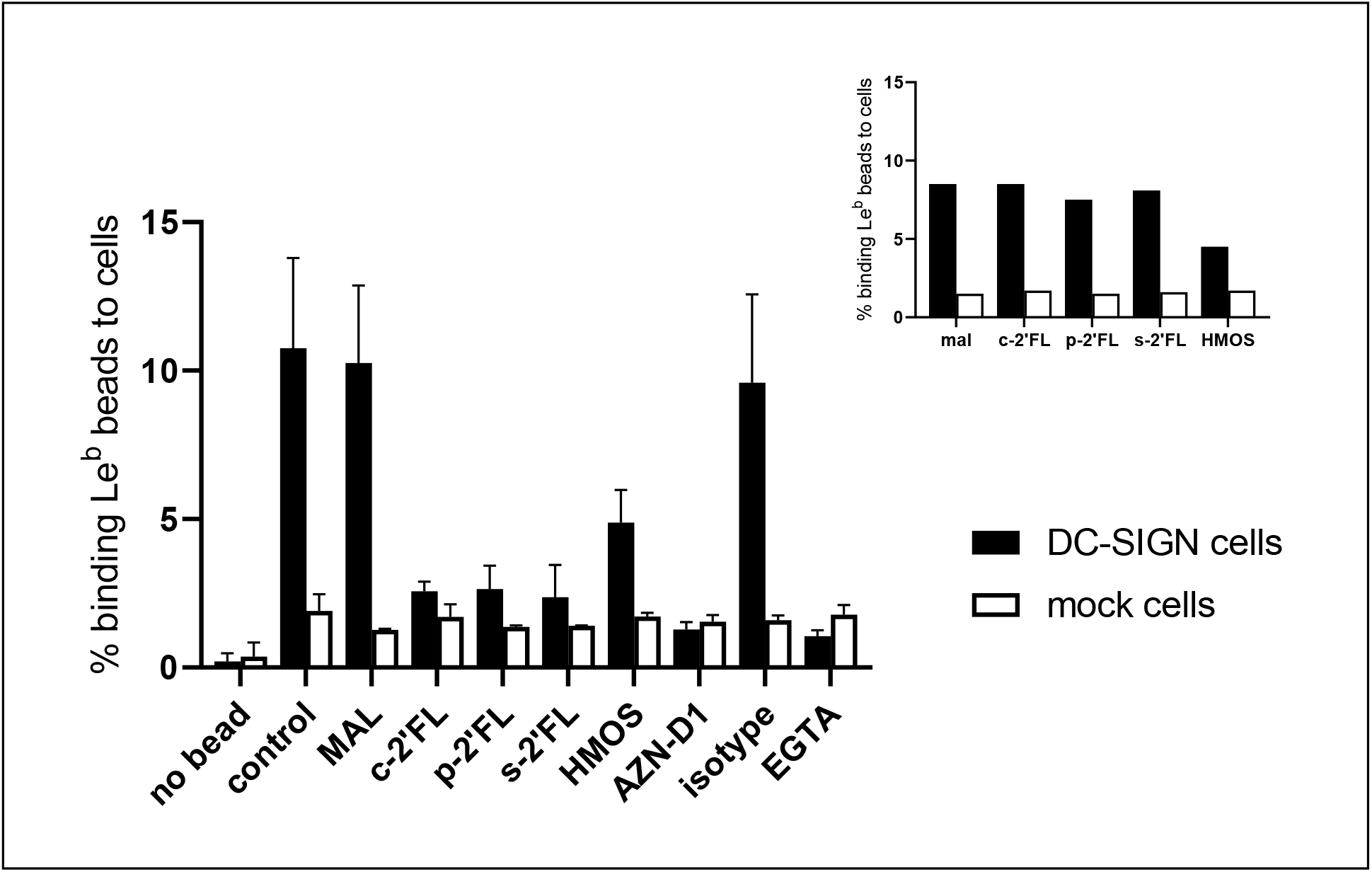
Cell-ligand competition assay with DC-SIGN expressing cells and mock cells and c2′-FL, p2′-FL, s2′-FL, HMOS, maltotriose (MAL, negative control of the 2′-FLs and HMOS), isotype antibody (negative control), AZN-D1 (DC-SIGN binding antibody), EGTA (as inhibition control). At a 5 mg/mL concentration of c2′-FL, p2′-FL, s2′-FL, HMOS inhibit the binding of Le^b^ coated fluorescent beads to DC-SIGN expressing cells. The insert shows the inhibition effect in 1 mg/mL concentration of c2′-FL, p2′-FL, s2′-FL, HMOS. Results are presented as mean % of cells binding Le^b^-coated fluorescent beads +/-SD, n=2. For insert, n=1. c-2′-FL = biotechnologically produced commercially available 2′-FL, p2′-FL = purified c-2′-FL and s2′-FL = chemically synthesized 2′-FL.

### Molecular Dynamics simulation suggested that 2′-FL exists in bioactive conformation to bind DC-SIGN but not Langerin

For the first time, an MD simulation study was performed to get more insight into the binding mode of DC-SIGN with 2′-FL. The simulation showed that the terminal fucose in 2′-FL acts both as a hydrogen bond donor and acceptor (Fig. 4). It remains hydrogen-bonded with DC-SIGN most of the time throughout the duration of the simulation (400 ns) (see Table S I for detailed interactions). The terminal fucose in 2′-FL remains hydrogen-bonded with DC-SIGN most of the time with glutamic acid (354 and 347) and aspartic acid (367) of DC-SIGN via hydrogen of -OH of fucose at 2, 4, and 3 positions respectively (as a hydrogen bond donor). The glucose anomeric -OH (as hydrogen bond donor) showed a connection to backbone oxygen and sidechain oxygen of serine (360) of DC-SIGN. Asparagine (365 and 349) of DC-SIGN was hydrogen-bonded via oxygen of -OH of fucose at 3 and 4 positions respectively (as hydrogen bond acceptor). All the hydrogen bonding interactions are shown in figure 4 and Table S I.

**Figure 4.**
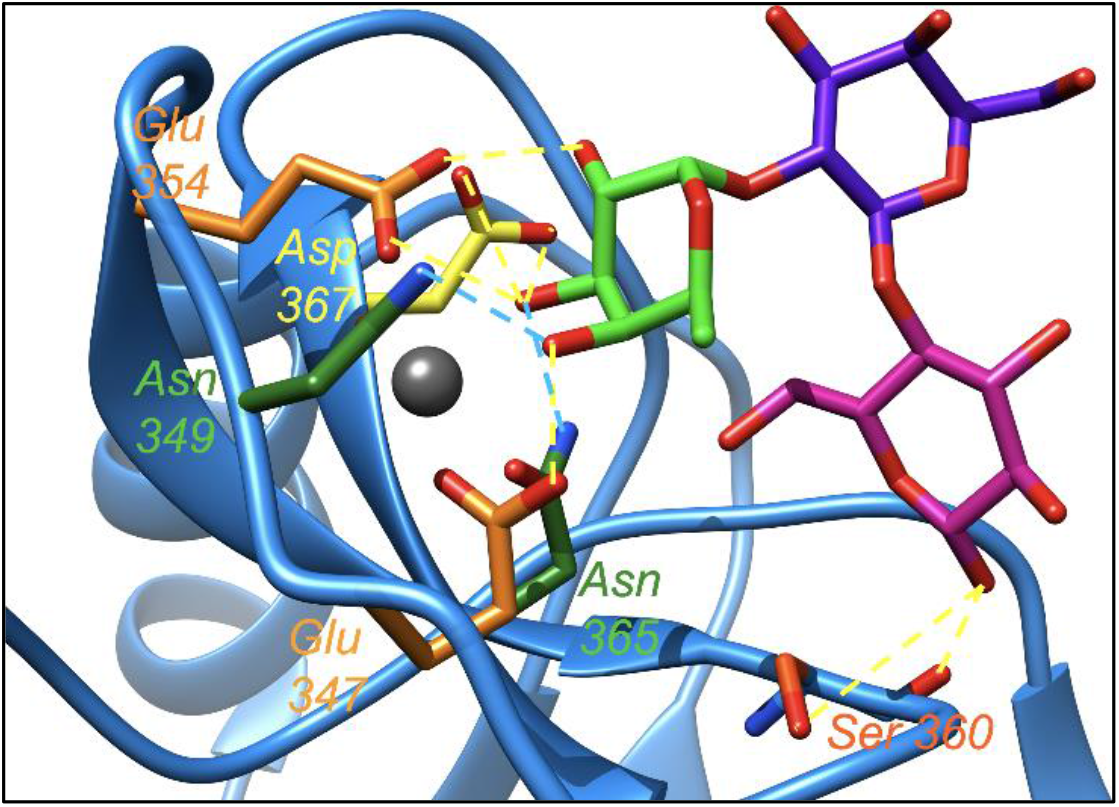
The hydrogen-bonding (dotted lines) network of 2′-FL and DC-SIGN. DC-SIGN interacting domain is represented as a blue ribbon. Fucose is coloured in green; galactose is coloured in dark blue; glucose is coloured in magenta; Ca^+2^ is coloured as a grey sphere. All the amino acids’ sidechain colours are matched with their respective label colours. The yellow dotted line represents interactions of 2′-FL as a hydrogen bond donor. The blue dotted line represents interactions of 2′-FL as a hydrogen bond acceptor.

The analysis of the binding mode and the dihedral angle of 2′-FL were performed to gain insight into its flexibility. 2′-FL remained quite rigid when it was bonded and non-bonded to DC-SIGN suggesting that 2′-FL preferentially displayed the bioactive conformation (pre-organized) before binding (Fig. S2) to DC-SIGN.

The binding affinity or stability of a complex depends on many factors. Non-covalent interactions, such as hydrogen bonds contribute to stabilizing the complex of protein and its ligand. Preorganized ligands also form the complex with its receptors with better affinity or stability due to its less entropic penalty (please see the section ‘Explanation of the possible better binding of 2′-FL to DC-SIGN’ in SI for the detailed explanation). As 2′-FL is preorganized and forms a hydrogen-bonding network with DC-SIGN, the affinity or stability of the complex of DC-SIGN and 2′-FL will be better than other flexible ligands. Using MMPBSA we calculated the enthalpy of the interaction of 2′-FL and DC-SIGN and that is 0.6 ± 0.4 Kcal/mol.

The MD simulation with 2′-FL and Langerin revealed that there was no binding between them. 2′-FL left the binding site of Langerin after 40 ns. When a ligand leaves the binding pocket, it means the molecule has lower energy (Gibbs) in the free state than in the bound state. Thus, the binding between the two entities may be very weak or none.

### Le^b^ interacts with DC-SIGN with two fucoses operating one at a time: consequences for displacement by 2′-FL

In Le^b^ one fucose is connected with galactose while the other fucose is connected with N-acetylglucosamine. The MD simulation showed that Le^b^ binds DC-SIGN with its two fucoses in two different ways. These two different binding modes appeared one at a time. When Le^b^ binds to DC-SIGN via its fucose connected to galactose, the following hydrogen-bonding networks were observed in MD simulation (see Table S II for detail). Most of the time glutamic acids (347 and 354) and aspartic acid (367) of DC-SIGN were connected to fucose (which is connected to galactose) via hydrogen of -OH of fucose at 4, 2 and 3 positions respectively (as hydrogen bond donor). Asparagine residues (349 and 365) of DC-SIGN were connected via oxygen of -OH of fucose at 4 and 3 positions respectively (as hydrogen bond acceptor). The hydrogen bonding connection between N-acetylglucosamine via O6 and arginine (345) of DC-SIGN was also observed. The fucose ring oxygen was also connected with asparagine (349). The interactions are shown in figure 5.

**Figure 5.**
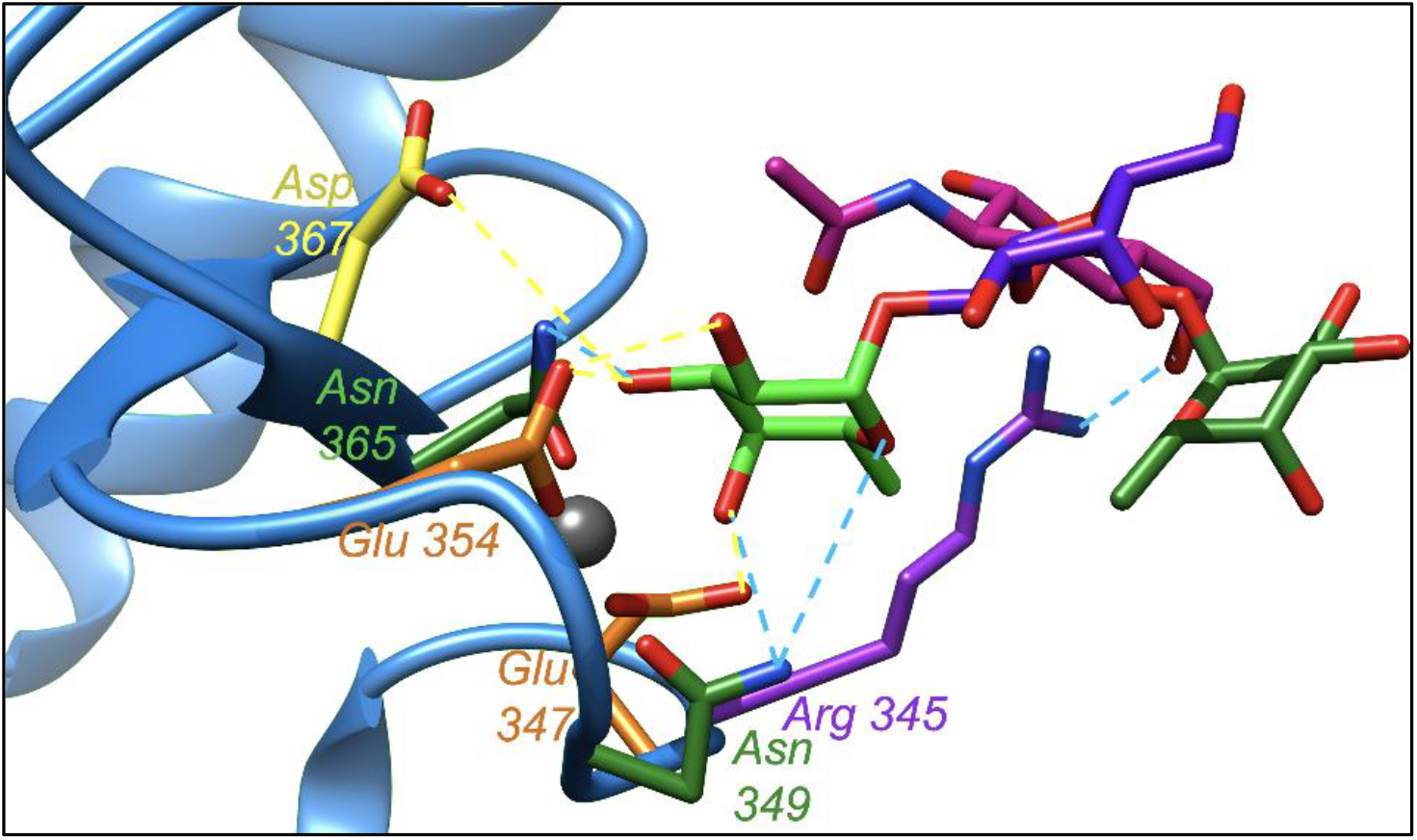
The hydrogen bonding (dotted lines) network of Le^b^ and DC-SIGN (bonded fucose is linked to the galactose). DC-SIGN interacting domain is represented as a blue ribbon. Fucose is coloured in green; galactose is coloured in dark blue; N-Acetylglucosamine is coloured in magenta; Ca^+2^ is coloured as a grey sphere. All the amino acids sidechain colours are matched with their respective label colours. The yellow dotted line represents interactions of Le^b^ as a hydrogen bond donor. The blue dotted line represents interactions of Le^b^ as a hydrogen bond acceptor.

The other binding mode of Le^b^ with the fucose which is connected to N-acetylglucosamine is described in Figure S3.

The MD trajectory analysis showed an important hydrogen-bond network established between Le^b^ and the DC-SIGN. These interactions were comparable with the interaction network found for the 2′-FL. However, Le^b^ displayed higher flexibility in water than 2’-FL in the free state and more than one conformation co-exists in the solution (Fig. S2). RMSD calculated over the heavy atoms along the 1 microsecond trajectory showed the coexistence of more than one conformation in the case of Le^B^ and just one conformation for the 2’-FL (Fig S4). This increased freedom of the dihedral angles contributes to augmenting the entropy penalty which may diminish the binding affinity of this ligand to DC-SIGN. Using MMPBSA we calculated the enthalpy of the interaction of Le^b^ and DC-SIGN. The enthalpy is 0.5 ± 0.4 Kcal/mol when Le^b^ is interacting with the fucose connected to galactose. The enthalpy is 7.8 ± 0.7 Kcal/mol when Le^b^ is interacting with fucose connected to GlcNAc.

The Ramachandran-like plot (Fig. S5) showed that 2′-FL adopted predominantly the bioactive conformation. However, Le^b^ showed restricted bioactive conformation in the majority in the bound state although a higher degree of flexibility than 2′-FL in the free state. The highly restricted conformation in the bound state for Le^b^ and its larger degree of flexibility in the free state cause a higher entropic penalty for binding to DC-SIGN than that of 2′-FL. Thus, Le^b^ may bind to DC-SIGN with a lower affinity than 2′-FL.

2′-FL shows similar binding enthalpy to Le^b^ when Le^b^ is interacting with fucose connected to galactose. There is a significant difference, however, in the enthalpy of interaction when Le^b^ is interacting with fucose connected to N-Acetylglucosamine. The lower binding enthalpy of 2′-FL suggests that it may bind to DC-SIGN with a higher affinity than that of Le^b^. As the two binding modes of Le^b^ operate one at a time, while switching the binding mode, there may be an opportunity for 2′-FL to enter into the binding pocket of DC-SIGN.

## Discussion

In line with previous findings, we have shown that 2′-FL binds to DC-SIGN but not to Langerin in the tested experimental conditions. In addition, our data show that in both ligand-receptor competition assays and cell-ligand competition assays, 2′-FL competes with the DC-SIGN ligands fucose monosaccharide and Le^b^ for binding to DC-SIGN in a dose-dependent way. It is worth mentioning that 2′-FL can compete with the DC-SIGN ligands fucose monosaccharide and Le^b^ even though these ligands were represented on PAA bead as multivalent ligands. Moreover, the observed inhibitory effect of 2′-FL on DC-SIGN ligand binding is comparable to that of a total HMOs mixture.

Recently, it has become possible to produce 2′-FL biotechnologically on a large scale (Bych et al. 2019). It is unclear whether 2′-FL immunomodulatory effects are due to the HMO itself or rather due to the presence of side products (Perdijk et al. 2018 in Glycobiology). Earlier work indicated that bacterial production of bioactive proteins such as heat shock proteins or other endogenous proteins might have immunomodulating effects linked to microbial contaminants rather than the actual protein (Erridge C. 2010). Similar effects are also published on the immune effects of HMOs whereupon rigorous purification of the HMOs, the immune effects were significantly reduced (Perdijk et al. 2018 in PLoS ONE). However, recent work by Sodhi et al (Sodhi et al. 2021), using experimental and modelling techniques showed that 2′-FL can bind to TLR4 and inhibits its activation by its ligand lipopolysaccharide (LPS) suggesting that 2′-FL might indeed directly bind to host receptors. Here we show that all forms of 2′-FL, either synthetically or biotechnologically produced as well as purified, can compete with fucose or Le^b^ for binding to DC-SIGN. Our results show that possible side products in biotechnologically produced 2’-FL are not playing a role in 2’-FL-induced inhibition of the binding of prototypical ligands to DC-SIGN.

There is a concentration difference in the effects of 2′-FL on the DC-SIGN-ligand interactions assessed by ligand-receptor and cell-ligand competitions assays. In the ligand-receptor competition assay, the effective concentration of all forms of 2′-FLs is 1mg/mL (2mM). This concentration is similar to previously reported, where it was also shown that IC_50_ of 2′-FL towards DC-SIGN binding is in the physiological concentration (1mM) (Noll et al. 2016). In addition, in human milk, the typical physiological concentration of 2′-FL is 0.5-2.5 g/L (1-5 mM) (Noll et al. 2016). In the cell-ligand competition assays, however, the concentration of 2′-FL needed to inhibit DC-SIGN-Le^b^ interaction is higher (10 mM).

Our MD simulation data supports the insight that 2′-FL directly binds to the CRD of DC-SIGN. 2′-FL remains mostly hydrogen-bonded with DC-SIGN during the entire simulation time (400ns). 2′-FL exists in bioactive conformation before binding to DC-SIGN, whereas Le^b^ has a flexible structure before binding to DC-SIGN. Moreover, 2′-FL and Le^b^ have comparable binding enthalpy when Le^b^ is interacting with fucose connected to galactose. However, Le^b^ has significantly higher binding enthalpy when Le^b^ is interacting with fucose connected to N-Acetylglucosamine. Due to the pre-existence of bioactive conformation, 2′-FL binds to DC-SIGN with a less entropic penalty and its lower binding enthalpy suggests that the binding affinity of 2′-FL to DC-SIGN may be stronger than that of Le^b^. Therefore, 2′-FL can inhibit the binding of Le^b^ to DC-SIGN. Also, Le^b^ has two binding modes, one via fucose connected to galactose and another via fucose connected to N-Acetylglucosamine. These binding modes are in action one at a time. This suggests that the Le^b^ may leave the binding pocket of DC-SIGN while changing its binding mode and making a room for other ligands like 2′-FL to bind. The presence of two fucoses does not add advantages to Le^b^ over 2′-FL to bind stronger to DC-SIGN. Le^b^ is not large enough to interact with binding pockets of two DC-SIGN molecules at the same time. Hence, strengthening the binding capacity of Le^b^ through multivalency is not possible.

The MD simulation data regarding the interaction between Langerin and 2′-FL demonstrated that 2′-FL leaves the binding pocket of Langerin after 40 ns suggesting that 2′-FL does not bind to Langerin. Although DC-SIGN and Langerin bind to their ligands using ‘EPN’ amino acid motif in CRDs, 2′-FL does not bind to Langerin suggesting that Langerin has a stronger affinity to its ligands or it has different requirements to bind to a fucosylated ligand. In general, the relative binding affinity of Lewis antigens to DC-SIGN-Fc is of the following orders: Le^b^ (47%)> Le^y^ (45%) >Le^a^ (34%)> Le^x^ (22%) (van Liempt et al 2006). In our study, we have demonstrated that 2′-FL replaces Le^b^ (presented on multivalent PAA constructs), which has the strongest affinity to DC-SIGN amongst the Lewis antigens (van Liempt et al 2006). Therefore, 2′-FL can replace other Lewis antigens or fucosylated ligands, which are more flexible than 2′-FL and have a similar or less binding affinity, from the CRD of DC-SIGN.

This study implicates that 2′-FL binds to DC-SIGN but not to Langerin and 2′-FL is able to outcompete Lewis antigens for binding to the CRD of DC-SIGN. Based on the importance of Lewis antigens in host-microbe interactions (Ewald et al. 2018) at the mucosal surface as well as the anatomical localization and expression of DC-SIGN on myeloid subsets of immune cells, these interactions might be of importance in shaping the microbiota and immune balance in health and disease (Triantis et al. 2018).

## Materials and methods

### Antibodies and beads

TSA buffer: 5ml 10x TSM + 0.5gr BSA (Millipore, 126575, Lot 3134664) + 45ml Milli-Q (ppb<2) BSA: (Sigma-Aldrich, lyophilized powder, ≥96%, agarose gel electrophoresis), HBSS: Gibco; EGTA, Sigma-Aldrich, E3889: ; AZN-D1: isolated as described in the ref Geijtenbeek et al. 2000 in *Cell, pp 575-585*: anti-human horseradish peroxidase: Jackson ; mouse IgG1 (isotype control): Invitrogen Lot 4347632, clone P3.6.2.8.1 cat 16-4714-82; 3,3′,5,5′-Tetramethylbenzidine, TMB, Sigma-Aldrich 860336 ; Streptavidin: Sigma S4762, Lot #098M4008V; Tween 20, Sigma-Aldrich, P1379: ; α-L-fucose-PAA beads and Le^b^-PAA beads: Lectinity, heck ; TransFluoSpheres carboxylate-modified microspheres: 488/645 nm, 1.0 μm; Molecular Probes, Eugene, OR.

### Cells

DC-SIGN expressing cells (OUW-SIGN cells) are generated by the transduction of EBV-BLCL with DC-SIGN as previously described (Marzi et al. 2007; Geijtenbeek et al. 2000, pp 587-97).

### HMOs

c2′-FL and whole HMOS were generously donated by Danone Nutricia, The Netherlands. p2′-FL was made by purifying c2′-FL. Synthetic 2′-FL (s2′-FL) was purchased from Carbosynth (www.carbosynth.com). The c2′-FL contains 94.1 % 2′-FL, 1.4% 3-FL, 1.5% DFL, 2.6% fucosyl-galactose, 0.1% fucose (measured by HAPEC-PAD). The s2′-FL contains 95% 2′-FL (purity min 95% by ^1^H NMR as mentioned by Carbosynth). The component analysis of HMOS is not done.

### Purification of c2′-FL to obtain p2′-FL

1 g of c2’-FL was dissolved in 10 mL of MilliQ water. The solution was poured into a separatory funnel. The solution was extracted with chloroform (3 × 5 mL). Then 300 mg of activated charcoal (pre-washed with MilliQ water) was added to the solution and left for 1 hr with occasional stirring. The charcoal was filtered with Whatman filter paper and finally with 2µM pore-sized filter paper. Then the water layer was lyophilized to collect 800 mg of p2′-FL (which contains mostly 2′-FL with a trace of 3-FL and DFL as observed in MALDI-MS).

### Ligand-receptor competition assay

The Polyacrylamide polymers (PAA) functionalized with different glycans were purchased from Lectinity, heck MW approx. 20 KDa, Carbohydrate content around 20% mol. 50 µL of Le^b^-PAA or Fucose-PAA at 1.0 µg/mL in PBS (10mM, pH=7.4) were used to coat the Nunc MaxiSorp plates overnight at room temperature. After discarding and washing with PBS (2×200µL), the wells were blocked with 200 µL of 1% BSA (Sigma-Aldrich, lyophilized powder, ≥96%, agarose gel electrophoresis) in HBSS containing CaCl_2_/MgCl_2_ (Gibco) at 37 degrees for 30 min. In parallel, DC-SIGN-Fc (5 µg/mL or 0.5 µg/mL or 0.1 µg/mL) or Langerin-Fc (5 µg/mL) was pre-incubated (competition) with serial dilutions of each c2′-FL p2′-FL, s2′-FL, and HMOS in assay buffer (1% BSA in HBSS), for 30 min at 37 degrees. Both chimeric constructs, DC-SIGN-Fc and Langerin-Fc, consist of their extracellular domains fused to the Fc portion of human IgG1. They were produced from established transfectants as described previously (Bloem et al. 2013; Fehres et al. 2015; Geijtenbeek et al. 2002).

The blocking solution was discarded from the plate wells, and the mixture of DC-SIGN-Fc (at concentrations of 5.0, 0.5 or 0.1 µg/mL) and each of c2′-FL, p2′-FL, s2′-FL, or HMOS (at concentrations of 1.0, 10, 100 and 1000 µg/mL) in assay buffer (1% BSA in HBSS) were added to the plate coated with Le^b^-PAA or Fucose-PAA.

After 2h at room temperature, the wells were washed with 0.05% Tween-20 in HBSS (3×200µL) and then 50 µL of anti-human horseradish peroxidase (0.5 µg/mL, Jackson, Goat anti-Human IgG-HRP) were added. After 30 min at room temperature, the wells were washed with 0.05% Tween-20 in HBSS (4×200µL).

Finally, 100 µL of substrate solution (3,3′,5,5′-Tetramethylbenzidine, TMB, in citric/acetate buffer, pH=4, and H_2_O_2_) were added and after 5 min incubation at room temperature the reaction was stopped with 50 µL of H_2_SO_4_ (0.8 M) and the optical density was measured at 450 nm in an ELISA reader. The experiment was performed in duplicate, and data were normalized over the signal at 450 nm from the non-coating wells. Data were expressed as % of binding and they were normalized over the OD signal from the wells without any competition. This OD value was set as 100% of interaction.

### Fluorescent bead adhesion cell-ligand competition assay

To demonstrate the effect of 2′-FLs and HMOS on the binding of Le^b^ to whole cells, a fluorescent beads adhesion assay was used as previously described (Geijtenbeek et al 1999). TransFluorSpheres were covalently coupled to Streptavidin. Then biotinylated PAA-Le^b^ beads were coupled to the streptavidin beads. Briefly, 7,5μl of streptavidin beads (+/-8.25 × 10^7^ beads) and 10μl solution of biotinylated PAA-Le^b^ (of 1mg/ml concentration) were taken in an Eppendorf and 300μl of PBA was added into that. The mixture was Incubated at 37 degrees for 2hrs, rotating at 550rpm. After incubation, the mixture was centrifuged at 13000 rpm using an Eppendorf centrifuge. The beads were washed 2x using 500μl PBA. The beads were suspended in 50μl PBA (+/-8.25 x107 beads/50μl).

DC-SIGN expressing cells (OUW-SIGN) and mock cells were made to a cell concentration of 10^6^ cells/ml in TSA. Then put 50μl/well of cells to a 96-well V-bottom plate (50.000 cells/well). Then each of c2′-FL, p2′-FL, s2′-FL, and HMOs were added either in 5mg/ml or 1mg/ml as the final concentration. EGTA (final concentration 10mM), isotype antibody (final concentration 25µg/mL), AZN-D1 (final concentration 25µg/mL) were added into the well. After addition of all the components, the plate was incubated for 15 minutes at 37°C. Then 10μl/well Le^b^-PAA-beads (final 1μl beads/well) were added. Then the plate was again incubated for 30 minutes at 37°C. FACS acquisition on Fortessa was performed immediately and cells were put at the ice. Beads were measured in channel Blue A-A at Voltage 400. In total 30,000 events were recorded.

### Molecular dynamics simulation

Interactions between the DC-SIGN and the different ligands, 2′-FL, Le^b^ were further evaluated by Molecular Dynamics simulations. In the case of Le^b^, two trajectories were run to analyze the recognition process, one trajectory for each of the two fucoses that this compound possesses. DC-SIGN and ligand complexes were built by using the crystal structure (PDB id: 1sl5) where DC-SIGN is complexed with the ligand LNFP III (fucosylated ligand). The LNFP III was replaced by each of our glycans. The complex was solvated with the water model TIP3P (Jorgensen et al 1983) in a 10 Å cubic box. The simulations were carried out by using the AMBER 18 suite (Case et al. 2018), the protein parameters were taken from ff14SB (Maier et al. 2015) force field and the glycan moiety from GLYCAM_06j-1 force field (Kirschner et al. 2008). In the case of Langerin, the complex was built by using the crystal structure of the Langerin RBD in a complex with blood group B trisaccharide (PDB id: 3P5G). Blood group B trisaccharide was replaced by 2′-FL prior to running the simulation. The protocol for the MD simulations applied to the molecules in the bound state starts with a two-step minimization process: first, the solute was fixed, and only solvent molecules were allowed to move; and during the second step, all the atoms were allowed to minimize in the simulation cell. Afterward, a three-step molecular dynamic simulation was performed, and the temperature was raised from 0 to 300 K under the constant pressure of 1 atm and periodic boundary conditions. Harmonic restraints of 10 kcal·mol^-1^ were applied to the solute, and the Berendsen temperature coupling scheme (Berendsen et al. 1984) was used to control and equalize the temperature. The time-step was kept at 1 fs during the heating stage. Water molecules were treated with the SHAKE algorithm such that the angle between the hydrogen atoms remained fixed. Particle-mesh-Ewald (PME) method was employed to model the long-range electrostatic interactions (Darden et al. 1993). An 8 Å cut-off was applied to Lennard-Jones and electrostatic interactions. Each system was equilibrated for 100 ps with a 2 fs timestep at a constant volume and temperature of 300 K. Production trajectories were then run for an additional 400 ns under the same simulation conditions. Protocol followed to perform MD simulation on the glycans in the free state is similar to the former one (bound state), but the production trajectory was extended up to 1 microsecond. Cpptraj module of AMBER 18 was employed to process the trajectory. MMPBSA was employed to calculate enthalpy values. 50 frames evenly separated among the last 5000 steps of the trajectory were considered for the enthalpy calculation.

## Funding

The work is supported by Nederlandse Organisatie voor Wetenschappelijk Onderzoek (Dutch Research council) LIFT grant (no. 731.017.408).

## Acknowledgements

Reshmi Mukherjee is thankful for the LIFT grant (no. 731.017.408). Victor J Somovilla is thankful for the financial support to EU Commission (Marie Skłodowska-Curie 840663 to V.J.S.) and Maria de Maeztu Units of Excellence Programme – Grant No. MDM-2017-0720 Ministry of Science, Innovation and Universities.

## Abbreviations

CRD: Carbohydrate recognition domain
CLR: C-type lectin receptors
DC-SIGN: Dendritic Cell-Specific Intercellular Adhesion Molecule-3-Grabbing Non-integrin
EBV-BLCL: Epstein-Barr Virus-transformed
B: lymphoblastoid cell line
ELISA: Enzyme-Linked Immunoassay
FACS: Fluorescence-activated cell sorting
Le^b^: Lewis B
HBSS: Hank’s balanced salt solution
HMOS: Human milk oligosaccharides
2′-FL: 2′-Fucosyllactose
EGTA: Egtazic acid
MD: Molecular Dynamics
PME: Particle-mesh-Ewald
mAbs: monoclonal antibodies
TMB: 3,3′,5,5′-Tetramethylbenzidine
TSM: Tris-Saline with Magnesium

## Data Availability Statement

The data underlying this article are available in the article and in its online supplementary material.

## Footnotes

Authors’ contribution: RM: Project concept and plan, purification of 2′-FL, data analysis, manuscript writing and communication and co-ordination between authors; VJS: MD simulations and critical comments; SB: Biological assays and data analysis; FC: critical comments, project plan and data analysis; RJP: critical comments; JG: valuable discussion; JvB: critical comments and manuscript writing; YvK: valuable discussion; AK: Data analysis and critical comments.

## Conflicts of Interests

JvB is a paid employee of Danone Nutricia Research. JG is partially paid by Danone Nutricia Research. There are no conflicts of interest by other authors of this manuscript.

## References

Asadpoor M, Peeters C, Henricks PAJ, Varasteh S, Pieters RJ, Folkerts G, Braber S. 2020. Anti-Pathogenic Functions of Non-Digestible Oligosaccharides In Vitro. Nutrients, 12, 1789.

Berendsen HJC, Postma JPM, van Gunsteren WF, DiNola A, Haak JR. 1984. Molecular dynamics with coupling to an external bath. J. Chem. Phys. 81, 3684–3690.

Bloem K, Garcia-Vallejo JJ, Vuist IM, Cobb BA, van Vliet SJ, van Kooyk, Y. 2013. Interaction of the capsular polysaccharide A from Bacteroides fragilis with DC-SIGN on human dendritic cells is necessary for its processing and presentation to T cells, Front. Immunol. 4: 103.

Bych K, Mikš MH, Johanson T, Hederos MJ, Vigsnæs LK, Becker P. 2019. Production of HMOs using microbial hosts— from cell engineering to large scale production. Curr. Opn. Biotechnol. 56. 130-137.

Case DA, Ben-Shalom IY, Brozell SR, Cerutt DS, Cheatham TE, Cruzeiro VWD, Darden TA, Duke RE, Ghoreishi D, Gilson MK, Gohlke H, Goetz AW, Greene D, Harris R, Homeyer N, Huang Y, Izadi S, Kovalenko A, Kurtzman T, Lee TS, LeGrand S, Li P, Lin C, Liu J, Luchko T, Luo R, Mermelstein DJ, Merz KM, Miao Y, Monard G, Nguyen C, Nguyen H, Omelyan I, Onufriev A, Pan F, Qi R, Roe DR, Roitberg A, Sagui C, Schott-Verdugo S, Shen J, Simmerling CL, Smith J, Salomon-Ferrer R, Swails J, Walker RC, Wang J, Wei H, Wolf RM, Wu X, Xiao L, York DM and Kollman PA. 2018. AMBER 2018, University of California, San Francisco. https://ambermd.org/

Cheng K, Zhou Y, Neelamegham S. 2017. ’DrawGlycan-SNFG: a robust tool to render glycans and glycopeptides with fragmentation information’, Glycobiology. 27: 200–205. http://www.virtualglycome.org/DrawGlycan/

Cheng L, Kiewiet MBG, Groeneveld A, Nauta A, de Vos P. 2019. Human milk oligosaccharides and its acid hydrolysate LNT2 show immunomodulatory effects via TLRs in a dose and structure-dependent way. J Functional Foods, 59, 174–184.

Chiodo F., de Haas A, van Vliet SJ, van Kooyk Y. 2021. Human C-Type Lectins, MGL, DC-SIGN and Langerin, Their Interactions with Endogenous and Exogenous Ligand Patterns. Comprehensive Glycoscience (Second Edition). 3; 425–441.

Darden T, Darrin Y, Pedersen L. 1993. Particle mesh Ewald: An N·log(N) method for Ewald sums in large systems. J. Chem. Phys. 98, 10089–10092.

Erridge C. 2010. Endogenous ligands of TLR2 and TLR4: agonists or assistants? J Leukoc Biol. 87, 989–999.

Ewald DR, Summer SCJ. 2018. Human Microbiota, Blood Group Antigens, and Disease. Wiley Interdiscip Rev Syst Biol Med. 10(3): e1413.

Fehres CM, Kalay H, Bruijns SCM, Musaafir SAM, Ambrosini M, van Bloois L, van Vliet S J, Storm G, Garcia-Vallejo JJ, van Kooyk Y. 2015. Cross-presentation through langerin and DC-SIGN targeting requires different formulations of glycan-modified antigens. Journal of Controlled Release. 203, 67–76.

Foster AJ, Bird JH, Timmer MSM, Stocker B. 2016. Pp 191-215. The Ligands of C-Type Lectins. In: Yamasaki S. (eds) C-Type Lectin Receptors in Immunity. Springer. Tokyo.

Geijtenbeek TB., Torensma, van Vliet SJ, van Duijnhoven G., Adema GJ., van Kooyk Y., Figdor CG. 2000. Identification of DC-SIGN, a novel dendritic cell-specific ICAM-3 receptor that supports primary immune responses. Cell. 100:575–585.

Geijtenbeek TB, Kwon DS, Torensma R, van Vliet SJ, van Duijnhoven GC, Middel J, Cornelissen IL, Nottet HS, KewalRamani VN, Littman DR, Figdor CG, van Kooyk Y. 2000. DC-SIGN, a dendritic cell-specific HIV-1-binding protein that enhances trans-infection of T cells. Cell. 100(5): 587–97.

Geijtenbeek TBH, van Duijnhoven GC, van Vliet SJ, Krieger E, Vriend G, Figdor CG, van Kooyk, Y. 2002. Identification of different binding sites in the dendritic cell-specific receptor DC-SIGN for intercellular adhesion molecule 3 and HIV-1. J. Biol. Chem. 277: 11314–11320.

Geijtenbeek, TBH, van Kooyk Y, van Vliet SJ, Renes MH, Raymakers RAP, Figdor C J. 1999. High Frequency of Adhesion Defects in B-Lineage Acute Lymphoblastic Leukemia. Blood. 94 (2): 754–764.

Goehring KC, Marriage BJ, Oliver JS, Wilder J.A., Barrett EG, Buck RH. 2016. Similar to those who are breastfed, infants fed a formula containing 2′-fucosyllactose have lower inflammatory cytokines in a randomized controlled trial. J Nutr Nutr Immunol. 146:2559–66.

He Y, Liu S, Kling D.E., Leone S, Lawlor N.T., et al. 2016. The human milk oligosaccharide 2′-fucosyllactose modulates CD14 expression in human enterocytes, thereby attenuating LPS-induced inflammation. Gut. 65:33–46.

Holscher HD, Bode L, Tappenden KA. 2017. Human Milk Oligosaccharides Influence Intestinal Epithelial Cell Maturation In Vitro. J Pediatr Gastroenterol Nutr, 64, 296–301.

Holscher HD, Davis SR, Tappenden KA. 2014. Human milk oligosaccharides influence maturation of human intestinal Caco-2Bbe and HT-29 cell lines. J Nutr. 144, 586–91.

Jorgensen WL, Chandrasekhar J, Madura JD, Impey RW, Klein ML. 1983. Comparison of simple potential functions for simulating liquid water. J. Chem. Phys. 79: 926–35.

Kirschner KN, Yongye AB, Tschampel SM, González-Outeiriño J, Daniels CR, Foley BL, Woods RJ. 2008. GLYCAM06: A Generalizable Biomolecular Force Field. Carbohydrates J. Comput. Chem. 29: 622–655.

Koromyslova A, Tripathi S, Morozov V, Schroten H, Hansman GS. 2017. Human norovirus inhibition by a human milk oligosaccharide. Virology. 508: 81–9.

Laucirica DR, Triantis V, Schoemaker R, Estes MK, Ramani S. 2017. Milk oligosaccharides inhibit human rotavirus infectivity in MA104 cells. J Nutr. 147:1709–14.

Lee RT, Hsu T-L, Huang SK, Hsieh S-L, Wong C-H, Lee YC. 2011. Survey of immune-related, mannose/fucose-binding C-type lectin receptors reveals widely divergent sugar-binding specificities. Glycobiology, 21, 512–520.

Maier JA, Martinez C, Kasavajhala K, Wickstrom L., Hauser KE, Simmerling C. 2015. ff14SB: Improving the Accuracy of Protein Side Chain and Backbone Parameters From ff99SB J. Chem. Theory Comput. 11(8):3696–3713.

Marzi A, Mӧller P, Hanna S. L., Harrer T, Eisemann J, Steinkasserer A, Becker S, Baribaud F, Pӧhlmann S, 2007. Analysis of the Interaction of Ebola Virus Glycoprotein with DC-SIGN (Dendritic Cell—Specific Intercellular Adhesion Molecule 3—Grabbing Nonintegrin) and Its Homologue DC-SIGNR. The Journal of Infectious Diseases. 196: S237–S246.

Merad M, Ginhoux F, Collin M. 2008. Origin, homeostasis and function of Langerhans cells and other langerin-expressing dendritic cells. Nature Reviews Immunology. 8:935–947.

Newburg DS. 2013. Glycobiology of human milk. Biochemistry (Mosc). 78:771–785.

Noll AJ, Yu, Y, Lasanajak, Y, Duska-McEwen, G, Buck, RH, Smith DF, and Cummings, RD. 2016. Human DC-SIGN Binds Specific Human Milk Glycans. Biochem J. 473(10): 1343–1353.

Pederson K, Mitchell DA, Prestegard JH. 2014. Structural Characterization of the DC-SIGN–LewisX Complex. 53 (35): 5700–5709.

Perdijk O, van Neerven RJJ, Meijer B, Savelkoul HFJ, Brugman S. 2018. Induction of human tolerogenic dendritic cells by 3′-sialyllactose via TLR4 is explained by LPS contamination. Glycobiology. 28:126–130.

Perdijk O, van Neerven J, van Den Brink E, Savelkoul HFJ, Brugman S. 2018. The oligosaccharides 6’-sialyllactose, 2’-fucosyllactose or galactooligosaccharides do not directly modulate human dendritic cell differentiation or maturation. PLoS ONE 13(7): e0200356

Ruiz-Palacios GM, Cervantes LE, Ramos P, Chavez-Munguia B, Newburg DS. 2003. Campylobacter jejuni binds intestinal H(O) antigen (Fuca1, 2Galb1,4GlcNAc), and fucosyloligosaccharides of human milk inhibit its binding and infection. J Biol Chem. 278:14112–20.

Sodhi CP, Wipf P, Yamaguchi Y, Fulton WB, Kovler M, Niño DF, Zhou Q, Banfield E, Werts AD, Ladd MR, Buck RH, Goehring KC, Prindle Jr T, Wang S, Jia H, Lu P, and Hackam DJ. 2021. The human milk oligosaccharides 2’-Fucosyllactose and 6’-Sialyllactose protect against the development of necrotizing enterocolitis by inhibiting Toll-Like Receptor 4 signaling. Pediatr Res. 89, 91–101.

Thurl S, Munzert M, Boehm G, Matthews C, Stahl, B. 2017. Systematic review of the concentrations of oligosaccharides in human milk. Nutr Rev. 75(11): 920–933.

Triantis V, Bode L, Joost van Neerven RJ. 2018. Immunological Effects of Human Milk Oligosaccharides. Front Pediatr. 6, 190.

Urashima T, Asakuma S, Leo F, Fukuda K, Messer M, Oftedal OT. 2012. The predominance of type I oligosaccharides is a feature specific to human breast milk. Adv Nutr. 3:473S–482S.

Valverde P, Delgado S, Martinez JD, Vendeville JP, Malassis J, Linclau B, Reichardt NC, Cañada FJ, Jiménez-Barbero, Ardá A. 2019. Molecular Insights into DC-SIGN Binding to Self-Antigens: The Interaction with the Blood Group A/B Antigens. ACS Chem. Biol. 14 (7), 1660–1671.

Valverde P, Martznez JD, Cañada FJ, Ardá A, Jiménez-Barbero J. 2020. Molecular Recognition in C-Type Lectins: The Cases of DC-SIGN, Langerin, MGL, and L-Sectin. ChemBioChem. 21: 2999 –3025.

van Liempt E, Bank CMC, Mehta P, Garciá-Vallejo JJ, Kawar ZS, Geyer R, Alvarez RA, Cummings RD, van Kooyk Y, van Die I. 2006. Specificity of DC-SIGN for mannose- and fucose-containing glycans. FEBS Lett. 580, 6123–31.

van Vliet SJ, García-Vallejo JJ, van Kooyk YY. 2008. Dendritic cells and C-type lectin receptors: coupling innate to adaptive immune responses. Immunology and Cell Biology. 86, 580–587.

Weichert S, Jennewein S, Hüfner E, Weiss C, Borkowski J, Putze J, et al. 2013. Bioengineered 2′-fucosyllactose and 3-fucosyllactose inhibit the adhesion of Pseudomonas aeruginosa and enteric pathogens to human intestinal and respiratory cell lines. Nutr Res. 33:831–8.

Weis WI, Taylor ME, Drickamer K. 1998. The C-type lectin superfamily in the immune system. Immunol Rev. 163:19–34.

