## Supplemental Figure S1, S2, S3, S4, S5, Table I, Table II for "Human Milk Oligosaccharide 2’-fucosyllactose inhibits ligand binding to C-type lectin DC-SIGN but not to Langerin"

[a] shared last author

Figure S1: ELISA type competition assay for DC-SIGN-Fc binding to PAA-fucose or PAA-$Le^b$ with maltotriose (MAL, negative control), c2'-FL, p2'-FL, s2'-FL and HMOS [in various concentration of 1µg/mL, 10µg/mL, 100µg/mL and 1000µg/mL ]. EGTA [10nM] was used to demomstrate $Ca^{+2}$and $Mg^{+2}$-dependent binding of DC-SIGN and Langerin to the PAA-fucose and PAA-$Le^b$ ligands. A) inhibition by 2'-FLs and HMOS of the binding of DC-SIGN-Fc (0.5 µg/mL) to the ligand PAA-fucose ; B) inhibition by 2'-FLs and HMOS of the binding of DC-SIGN-Fc (0.1 µg/mL) to the ligand PAA-fucose; C) inhibition of 2'-FLs and HMOS on the binding of DC-SIGN-Fc (0.5 µg/mL) to the ligand PAA-$Le^b$ ; D) inhibition of 2'-FLs and HMOS on the binding of DC-SIGN-Fc (0.1 µg/mL) to the ligand PAA-$Le^b$ . Results are presented as mean % binding of ligand to DC-SIGN-Fc and Langerin-Fc +/- SD, n=2. No block condition is regarded as 100% ligand-DC-SIGN-Fc/Langerin-Fc binding (not shown in the graph). c-2'-FL = biotechnologically produced commercially available 2'-FL, p2'-FL = purified c-2'-FL and s2'-FL = chemically synthesized 2'-FL.

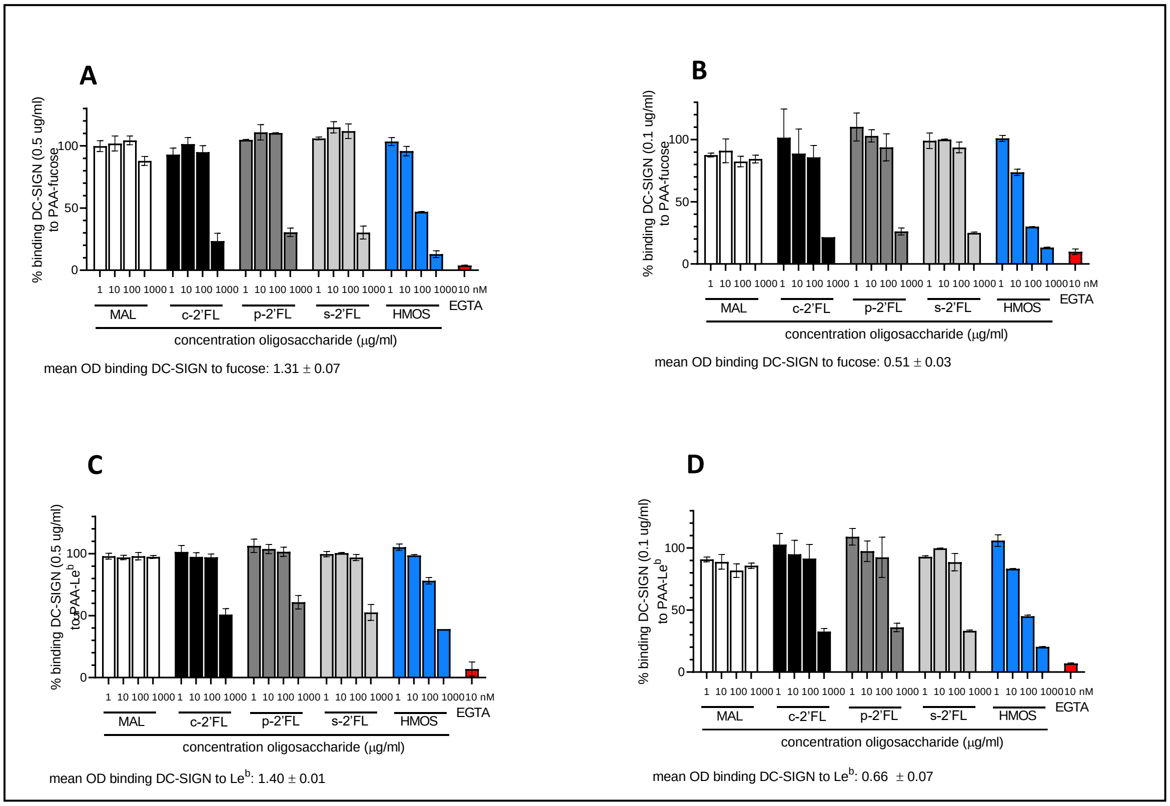

Figure S2: Overlay of conformational ensembles derived from MD simulations. Frames extracted from the MD trajectory of the glycans 2'-FL and Lewis B ($Le^b$) in the free state and the bound state in explicit water. 2'-FL shows one major conformation whereas Lewis B ($Le^b$) displays more flexibility in the free state. In the bound state both ligands present a similar behavior. Fucose is coloured in green, galactose is coloured in blue, glucose is coloured in magenta (for 2'-FL), N-acetylglucosamine is coloured in magenta (for $Le^b$).

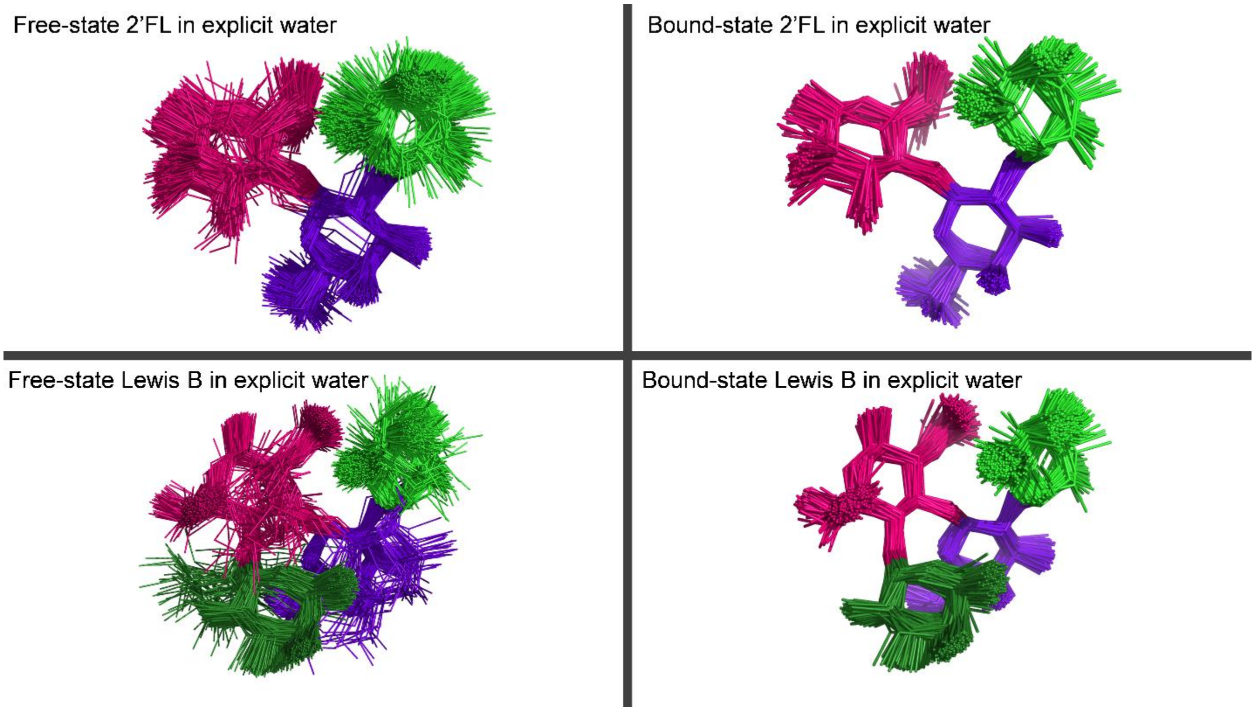

Figure S3: The other binding mode of $Le^b$ (with the fucose which is connected to N-acetylglucosamine) with DC-SIGN. DC-SIGN interacting domain is represented as a blue ribbon. Fucose is coloured in green; galactose is coloured in dark blue; N-acetylglucosamine is coloured in magenta; $Ca^{+2}$ is coloured as a grey sphere. All the amino acids sidechain colours are matched with their respective label colours. The yellow dotted line represents interactions of $Le^b$ as a hydrogen bond donor. The blue dotted line represents interactions of $Le^b$ as a hydrogen bond acceptor.

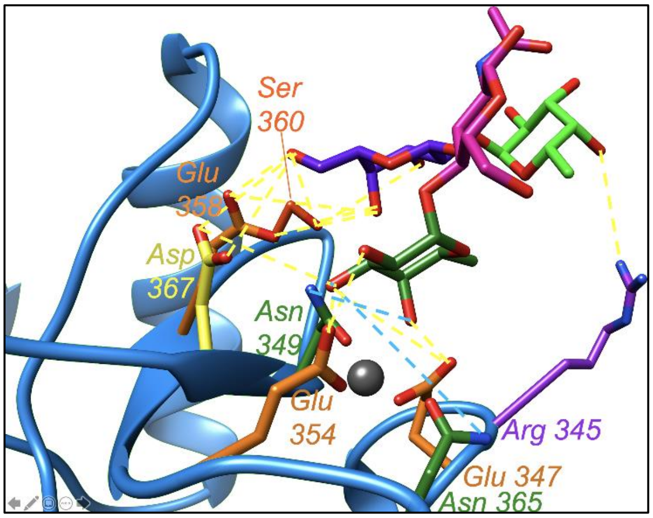


The MD simulation revealed that most of the time glutamic acids (347 and 354) and aspartic acid (367) of DC-SIGN were connected to fucose (which is connected to N-acetylglucosamine) via its hydrogen of the -OH of fucose at 4, 2 and 3 positions respectively (as hydrogen bond donor). The hydrogen bonding interactions was also observed between galactose via its O4 and O6 to two sidechain oxygens of glutamic acid (358) of DC-SIGN. The galactose O6 was also connected to two oxygens of aspartic acid (367). The galactose O3 was also connected to serine (360). Asparagine residues (349 and 365) of DC-SIGN were connected to fucose via oxygen of the -OH of fucose at 4 and 3 positions respectively (as hydrogen bond acceptor). The hydrogen bonding connections between galactose via O6 and serine (360) of DC-SIGN; fucose via O4 and arginine (345) of DC-SIGN were also observed.

Figure S4: RMSD calculated over the heavy atoms along the 1 microsecond trajectory.

RMSD plot against the time shows a major conformation present during the trajectory in both cases. There are two conformational changes for $Le^b$ in the free state , however, only one conformation is present for the 2'-FL glycan.

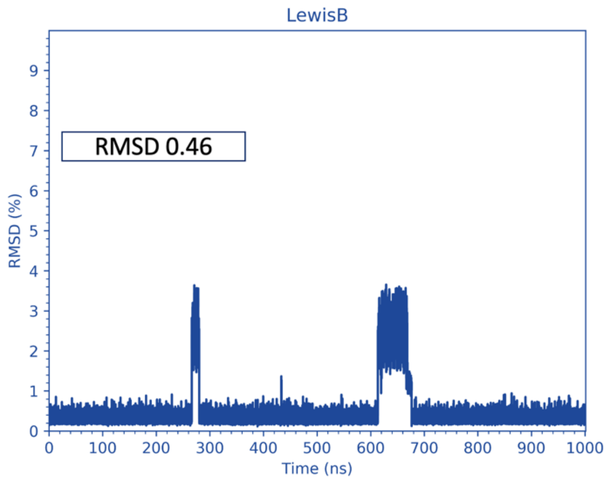


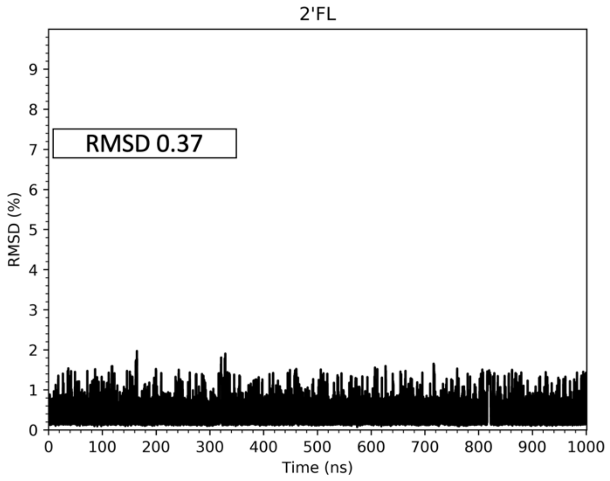

Figure S5: Ramachandran-like plots for the different glycosidic linkages

The following figures show the dihedral angles distribution for each glycosidic linkage both in the free- (purple) and the bound-state (yellow/blue). 2'-FL, in the bound state (yellow), shows flexibility for the bioactive conformation with two major populations for the linkage between the fucose and the galactose (glycosidic bond 2). Having a look at the free-state, we observe that 2'-FL adopts predominantly the bioactive conformation.

However, Lewis B (Le$^{b}$) displays a constrained conformation in the bound state, either when is recognized by the fucose linked to the galactose (yellow) or when interacts through the fucose linked to the N-acetylglucosamine (blue). Lewis B shows the bioactive conformation in majority although a higher degree of flexibility than 2'-FL in the free state. Thus, Lewis B entropy penalty for binding is higher than that of 2'-FL, due to the highly restricted conformation in the bound state for Lewis B and its larger degree of flexibility in the free-state.

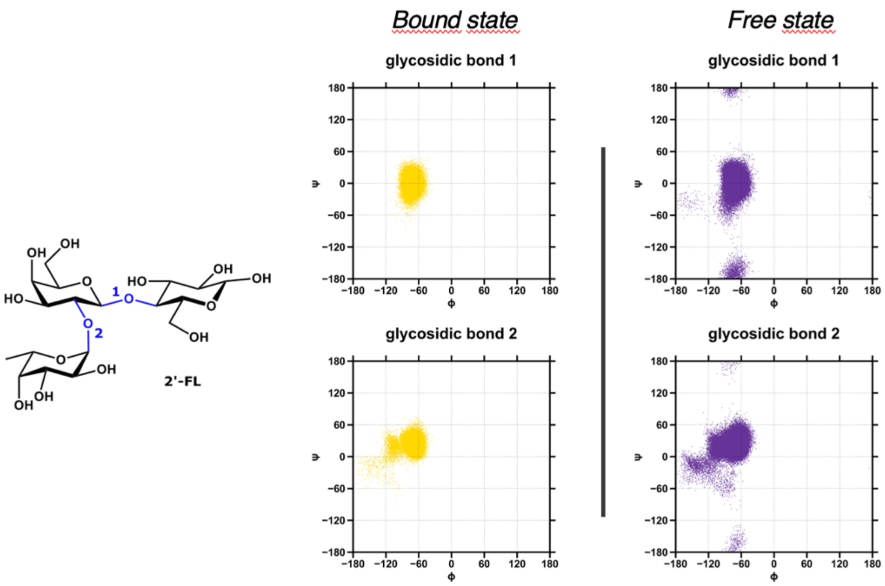

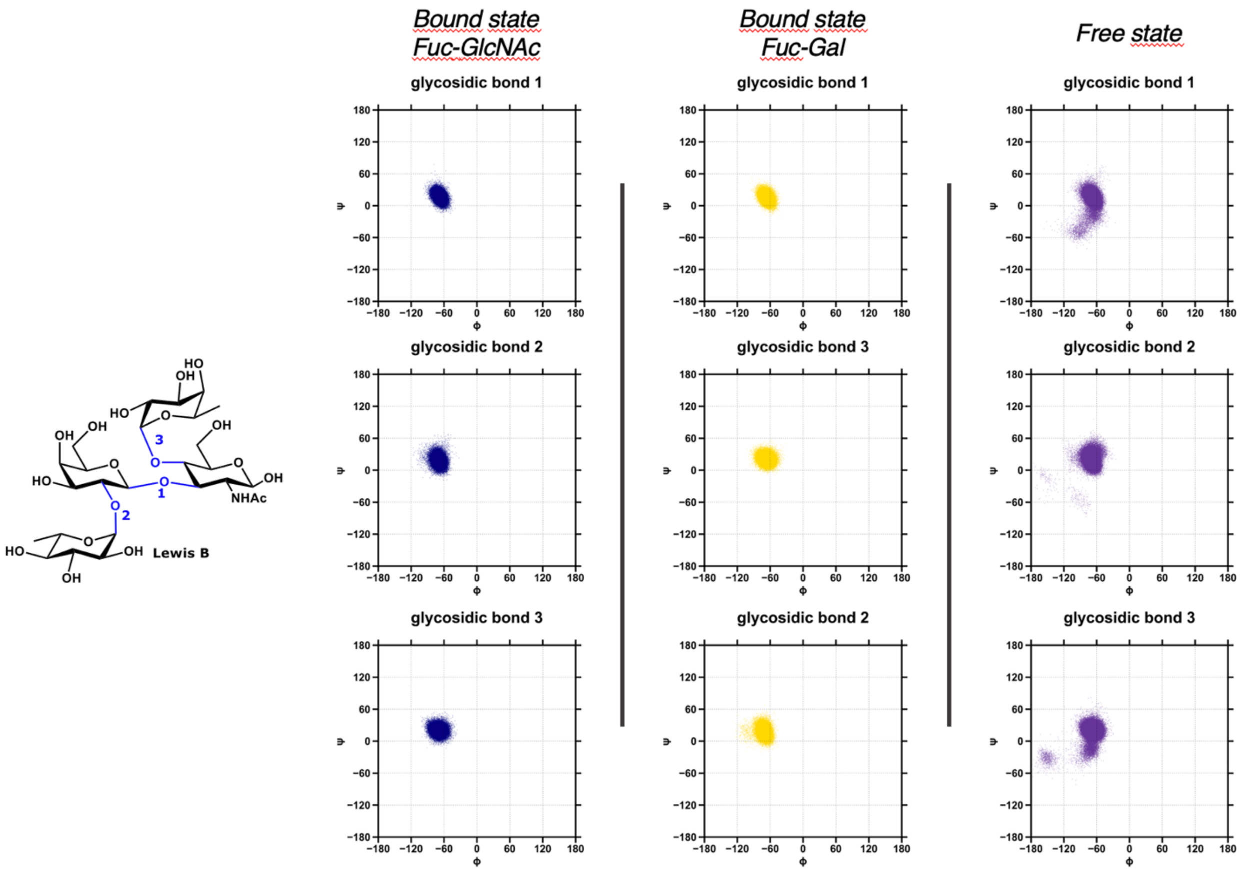

Bound state Fuc-GlcNAc
Bound state Fuc-Gal
Free state
glycosidic bond 1
glycosidic bond 2
glycosidic bond 3
glycosidic bond 1
glycosidic bond 3
glycosidic bond 2
glycosidic bond 1
glycosidic bond 2
glycosidic bond 3
ψ
φ
Lewis B
NHAc

Table S I: The hydrogen bonding networks of 2'-FL with DC-SIGN

2'-FL as a Hydrogen bond donor:

Glutamic (OE2) 354 – Fucose (O2) 99%;

Glutamic (OE2) 347 – Fucose (O4) 98%;

Aspartic (OD2) 367 – Fucose (O3) 40%;

Aspartic (OD1) 367 – Fucose (O3) 27%;

Glutamic (OE2) 354 – Fucose (O3) 17%;

Serine 360 (O) – Glucose (O1) 4%;

Serine 360 (OG) – Glucose (O1) 1%.

2'-FL as a Hydrogen bond acceptor:

Fucose (O3) – Asparagine 365 (ND2) 37%;

Fucose (O4) – Asparagine 349 (ND2) 4%

Table S II: The hydrogen bonding networks of Lewis B ($Le^b$) with DC-SIGN:

Fucose-Galactose (Interactions through Calcium ion and Fucose (390) linked to galactose (389)) as Hydrogen bond donor:

Glutamic 347 (OE2) – Fucose 390 (O4) 99%;

Glutamic 354 (OE2) – Fucose 390 (O2) 99%;

Glutamic 354 (OE2) – Fucose 390 (O3) 82%;

Aspartic 367 (OD2) – Fucose 390 (O3) 1%.

When the ligand as a Hydrogen acceptor:

Fucose 390 (O4) – Asparagine 349 (ND2) 31%;

Fucose 390 (O3) – Asparagine 365 (ND2) 20%;

N-Acetyl-Glucosamine (O6) – Arginine 345 (NH2) 1%;

Fucose 390 (O5) – Asparagine 349 (ND2) 1%.

Fucose-GlcNAc (Interactions through Calcium ion and Fucose (391) linked to N-acetyl glucosamine (388)) as Hydrogen bond donor:

Glutamic 354 (OE2) – Fucose 391 (O2) 98%;

Glutamic 347 (OE2) – Fucose 391 (O4) 97%;

Glutamic 354 (OE2) – Fucose 391 (O3) 36%;

Glutamic 358 (OE1) – Galactose 389 (O6) 20%;

Glutamic 358 (OE2) – Galactose 389 (O4) 9%;

Aspartic 367 (OD1) – Fucose 391(O3) 9%;

Aspartic 367 (OD1) – Galactose 389 (O6) 8%;

Aspartic 367 (OD2) – Galactose 389 (O6) 5%;

Glutamic 358 (OE1) – Galactose 389 (O4) 2%;

Serine 360 (OG) – Galactose 389 (O3) 1%;

Glutamic 347 (OE2) – Fucose 391 (O3) 1%;

Serine 360 (OG) – Galactose 389 (O4) 1%.

When the ligand as a Hydrogen bond acceptor:

Fucose 391 (O3) – Asparagine 365 (ND2) 36%;

Galactose 389 (O6) – Serine 360 (OG) 11%;

Fucose 391 (O4) – Asparagine 349 (ND2) 8%;

Fucose 390 (O4) – Arginine 345 (NH2) 1%.

**Explanation of the possible better binding of 2'-FL to DC-SIGN:**

The binding affinity or stability of a complex depends on the Gibbs free energy of the complex. The Gibbs free energy is dependent on enthalpy and entropy as evident from the equation $\Delta G = \Delta H - T\Delta S$, where $\Delta G$ is the change in Gibbs free energy (also known as binding free energy), $\Delta H$ and $\Delta S$ are the change in enthalpy and entropy of the system upon ligand binding respectively, T is the temperature in Kelvin. The protein-ligand binding occurs spontaneously only when $\Delta G$ of the system is negative after reaching equilibrium. The magnitude of the negative $\Delta G$ determines the stability of any protein-ligand complex, thus the binding affinity between the protein and its ligand.

Van der Waals interactions, hydrogen bonds, and electrostatic interactions contribute to stabilizing the complex of protein and its ligand by lowering its $\Delta G$ by decreasing its $\Delta H$. The $\Delta S$ contributes reversely to the binding affinity due to the minus sign in front of it. The higher the $\Delta S$ the lower the free energy, thus better the binding. Molecules with bigger flexibility (conformational and/or vibrational) will have a large negative $\Delta S$ value upon binding and as a result of this their binding affinity will be poorer. Alternatively, preorganized ligands will have a low negative or positive $\Delta S$ favouring the complex formation with better affinity or stability. As 2'-FL is preorganized, then the entropic penalty will be less compared to other flexible ligands. Hence, the binding free energy, thus affinity or stability of the complex of DC-SIGN and 2'-FL will be better than other flexible ligands such as $Le^b$.
